# WWP1 deficiency protects from cardiac remodeling induced by simulated microgravity

**DOI:** 10.1101/2021.06.18.447041

**Authors:** Guohui Zhong, Dingsheng Zhao, Jianwei Li, Zifan Liu, Junjie Pan, Xinxin Yuan, Wenjuan Xing, Yinglong Zhao, Shukuan Ling, Yingxian Li

## Abstract

Cardiac muscle is extremely sensitive to changes in loading conditions, the microgravity during space flight can cause cardiac remodeling and function decline. At present, the mechanism of microgravity-induced cardiac remodeling remains to be revealed. WW domain-containing E3 ubiquitin protein ligase 1 (WWP1) is an important activator of pressure-overload induced cardiac remodeling by stabilizing disheveled segment polarity proteins 2 (DVL2) and activating CaMKII/HDAC4/MEF2C axis. However, the role of WWP1 in the cardiac remodeling induced by microgravity is unknown. The purpose of this study was to determine whether WWP1 was also involved in the regulation of cardiac remodeling caused by microgravity. Firstly, we detected the expression of WWP1 and DVL2 in the heart from mice and monkeys after simulated microgravity using western blotting and Immunohistochemistry. Secondly, WWP1 knockout (KO) and wild type mice were subjected to hindlimb unloading (HU) to simulate microgravity effect. We assessed the cardiac remodeling in morphology and function through histological analysis and echocardiography. Finally, we detected the phosphorylation level of CaMKII and HDAC4 in the heart from WT and WWP1 KO mice after HU. The results revealed the increased expression of WWP1 and DVL2 in the heart both from mice and monkey after simulated microgravity. WWP1 deficiency protected against simulated microgravity-induced cardiac atrophy and function decline. Histological analysis demonstrated WWP1 KO inhibited the decreases in the size of individual cardiomyocytes of mice after hindlimb unloading. WWP1 KO can inhibit the activation of DVL2/CaMKII/HDAC4 pathway in heart of mice induced by simulated microgravity. These results demonstrated WWP1 as a potential therapeutic target for cardiac remodeling and function decline induced by simulated microgravity.

## Introduction

Our organ systems have evolved to work under 1g. It is not yet clear what effects will be produced by long-term exposure to low-gravity environments, and how these effects will be manifested at the cellular and molecular levels^[1]^. For many years, cardiac health has been a major concern of the world’s space agencies, however, the clinical evidence for decreased function and unmasking of asymptomatic cardiovascular disease is limited.

Cardiac muscle is extremely sensitive to changes in loading conditions^[2, 3]^. When exposed to microgravity during space flight, there are various changes in cardiac structure and function^[4, 5]^. Microgravity can cause a chronic decrease in metabolic demand and oxygen uptake, thereby reducing the demand for cardiac output, leading to cardiac atrophy and decline in cardiac function^[6, 7]^ There are many important factors that can regulate cardiac remodeling caused by various stimuli, including DVL2, CaMKII, and HDAC4^[8, 9]^. Our previous studies have shown that pathological cardiac remodeling signals (such as HDAC4) were activated in heart of mice after hindlimb unloading, which may lead to cardiac remodeling and decline in function^[3, 6]^. Walls et al. found that exposure to microgravity aboard the ISS caused heart dysfunction in a fly cardiac model, hearts are less contractile and exhibit changes in genes and proteins that maintain heart structure and function, among which, it is worth noting that proteasome genes expression was upregulated, suggesting that the regulation of protein homeostasis mediated by ubiquitination modification plays an important role in myocardial remodeling caused by microgravity^[1]^.

WW domain-containing E3 ubiquitin protein ligase 1 (WWP1) is a C2-WW-HECT-type ubiquitin E3 ligase containing an N-terminal C2 domain, four tandem WW domains and a C-terminal catalytic HECT domain for ubiquitin transferring^[10]^. WWP1 regulates a variety of cellular biological processes including protein trafficking and degradation, transcription and signaling by functioning as the E3 ligase^[9, 11–14]^. Cardiomyocyte-specific overexpression of WWP1 is detrimental to heart for it can induce arrhythmia and hypertrophy^[15]^. Also, in our previous research, we identified WWP1 as a novel regulator of pathological cardiac hypertrophy and heart failure. Pressure overload-induced heart hypertrophy was relieved in WWP1-KO mice. Mechanically, WWP1 stabilized DVL2 by promoting its K27-linked polyubiquitination, and regulated cardiac remodeling through DVL2/CaMKII/HDAC4/MEF2C pathway^[9]^. However, the role of WWP1 in cardiac remodeling induced by microgravity was unknown.

In this study, we found WWP1 protein levels was increased in the heart of mice and rhesus monkeys after simulated microgravity. WWP1 deficiency protected against simulated microgravity induced cardiac atrophy, function decline and pathological signals activation in mice, and WWP1 has potential as a therapeutic target for cardiac remodeling and function decline induced by simulated microgravity.

## Methods

### Animal experiments

The experimental procedures in mice and protocol used in this study were approved by the Animal Care and Use Committee of China Astronaut Research and Training Center. All animal studies were performed according to approved guidelines for the use and care of live animals (Guideline on Administration of Laboratory Animals released in1988 and 2006, Guideline on Humane Treatment of Laboratory Animals from China, and also referring to European Union guideline 2010/63).

The healthy rhesus monkeys with body weight of 5–8 kg and 4–8 years old were purchased from Beijing Xieerxin Biology Resource (Beijing, China). The monkeys were maintained at-10-degree head-down tilt position for 6 weeks. The whole process were supervised and monitored 24 h/day.

WWP1 knockout mice were bought from Model Animal Research Center of Nanjing University, as is reported previously (Zhao et al, 2021). All WT and WWP1 knockout (KO) mice used in this study were bred and housed at the specific-pathogen-free (SPF) Animal Research Center of China Astronaut Research and Training Center (12:12-h light-dark cycle, temperature: 23°C). All the experiments were repeated three times and were performed with homozygous WWP1 deficient mice (3 months old) and age-matched wild-type (WT) littermates. The hindlimb unloading procedure were elevated by tail suspension, as described before^[6]^. Briefly, the 3 months old mice were maintained in individual cage and suspended with a strip attached the tail and linked a chain hanging from a pulley. The mice were elevated to an angle of 30° to the floor with only the forelimbs touching the floor, which allowed the mice to move and access to food and water freely. The mice were retained to hindlimb unloading by tail suspension for 6 weeks, the height of hindlimb suspension was modulated to prevent the hindlimb from touching the ground. Control mice of the same strain background and the age-matched littermates and were instrumented and monitored in the identical cage conditions without tail suspension.

### Echocardiography

As described in our previous studies^[8]^, animals were lightly anesthetized with 2,2,2-tribromoethanol (0.2 ml/10 g body weight of a 1.2% solution) and set in a supine position. Two dimensional (2D) guided M-mode echocardiography was performed using a high-resolution imaging system (Vevo 1100, Visual-Sonics Inc., Toronto, Canada). Two-dimensional images are recorded in parasternal long- and short-axis projections with guided M-mode recordings at the midventricular level in both views. Left ventricular (LV) cavity size and wall thickness are measured in at least three beats from each projection. Averaged LV wall thickness [anterior wall (AW) and posterior wall (PW) thickness] and internal dimensions at diastole and systole (LVIDd and LVIDs, respectively) are measured. LV fractional shortening [(LVIDd – LVIDs)/LVIDd] and LV mass (LV Mass = 1.053 × [(LVID;d + LVPW;d + LVAW;d)^3^ – LVID;d^3^]) are calculated from the M-mode measurements. LV ejection fraction (EF) was calculated from the LV cross-sectional area (2-D short-axis view) using the equation LV %EF= (LV Vol;d – LV Vol;s) / LV Vol;d×100%. The studies and analysis were performed blinded as to experimental groups.

### RNA extraction and Real-time PCR

Total RNA was extracted from heart tissues with Trizol reagent according to the manufacturer’s protocol. The RNA (1 ug/sample) was reverse transcribed into cDNA and Q-PCR was performed using a SYBR Green PCR kit (Takara) in a Light Cycler (Eppendorf). The expression level of each gene was normalized to that of Gapdh, which served as an endogenous internal control. Primers (synthesized by BGI, China) for *BNP, Col1a1, Col3a1* and *Gapdh* were as follows:

*BNP* forward primer: 5’-TGTTTCTGCTTTTCCTTTATCTG-3’,
*BNP* reverse primer: 5’-TCTTTTTGGGTGTTCTTTTGTGA-3’;
*Col1a1* forward primer: 5’-CTGACTGGAAGAGCGGAGAGT-3’,
*Col1a1* reverse primer: 5’-AGACGGCTGAGTAGGGAACAC-3’;
*Col3a1* forward primer: 5’-ACGTAAGCACTGGTGGACAG-3’,
*Col3a1* reverse primer: 5’-CAGGAGGGCCATAGCTGAAC-3’;
*Gapdh* forward primer: 5’-ACTCCACTCACGGCAAATTCA-3’,
*Gapdh* reverse primer: 5’-GGCCTCACCCCATTTGATG-3’.

### Protein extraction and western blot

Heart tissues from mice or rhesus monkeys were crushed by homogenizer (Power Gen125, Fisher Scientific) and then lysed in lysis buffer (50 mM Tris, pH 7.5, 250 mM NaCl, 0.1% sodium dodecyl sulfate, 2 mM dithiothreitol, 0.5% NP-40, 1 mM PMSF and protease inhibitor cocktail) on ice for 30 min. Protein fractions were collected by centrifugation at 13,000 g at 4 °C for 15 min. Protein samples were separated by 10% SDS–PAGE and transferred to nitrocellulose membranes. The membranes were blocked with 5% bovine serum albumin and incubated with specific antibodies overnight. Antibodies used were: WWP1 (1:1000, Abcam, USA, #ab43791), DVL2 (1:1000, Cell Signaling Technology, USA, #3224S), CaMKII (1:1000, GeneTex, USA, GTX111401), p-CaMKII (1:1000, T287, GeneTex, USA, GTX52342), HDAC4 (1:1000, Cell Signaling Technology, USA, #5392), p-HDAC4 (1:1000, S246, Cell Signaling Technology, USA, #3443), GAPDH (1:2000, Abways Technology, China, #AB0036).

### Histological Analysis

Sections for Hematoxylin and eosin (H&E) and Masson’s Trichrome staining were generated from paraffin embedded hearts. Frozen sections were used to visualize cardiomyocyte cell membranes by staining with FITC-conjugated wheat-germ agglutinin (Sigma-Aldrich, USA), as described before Zhao et al. (2021). For immunohistochemical staining, sections were deparaffinized in xylene and rehydrated. Antigen retrieval was performed with protease K at 37 °C for 15 min. A solution of 3% H2O2 was used to block the activity of endogenous peroxidase. The sections were then incubated overnight at 4 °C with WWP1 (1/100, Abcam, #ab43791) or DVL2 antibody (1/100, Cell Signaling Technology, #3224S). After three times washes in PBS, biotinylated secondary antibodies were then added and incubated for 1 h at room temperature, followed by color development with DAB kit (ZSGB-bio). Negative control experiments were done by omitting the primary antibodies. The sections were examined using a microscope (ECLIPSE Ci-S, Nikon).

### Statistical Analysis

Data are presented as mean ± SEM. Statistical analysis for comparison of two groups was performed using two-tailed unpaired Student’s t-test. Statistical differences among groups were analyzed by 1-way analysis of variance (ANOVA) or 2-way ANOVA (if there were 2 factor levels), followed by Bonferroni’s post hoc test to determine group differences in the study parameters. Pearson correlation coefficients (r2) was performed to assess the correlation between two variables. All statistical analyses were performed with Prism software (GraphPad prism for windows, version 9.0, Nashville, USA) and SPSS (Version 20.0). Differences were considered significant at *P < 0.05, **P < 0.01, ***P < 0.001.

## Results

### Characterization of WWP1 expression during simulated microgravity induced cardiac remodeling

To assess potential role of WWP1 in cardiac remodeling induced by simulated microgravity, hearts from mice after 6 weeks of hindlimb unloading were assessed for WWP1 expression. As shown in **Figure 1A-C**, Western blotting and immunohistochemistry revealed that the protein level of WWP1 were significantly increased in the hearts of mice after hindlimb unloading. Moreover, we detected WWP1 expression in hearts of rhesus monkeys after 6 weeks of bed rest, and found that the protein levels of WWP1 increased in the heart of rhesus monkeys after bed rest (**Figure 1D-F**). These results suggested that simulated microgravity increase the protein level of WWP1, which was a activator of pathological cardiac remodeling.

**Figure 1.**
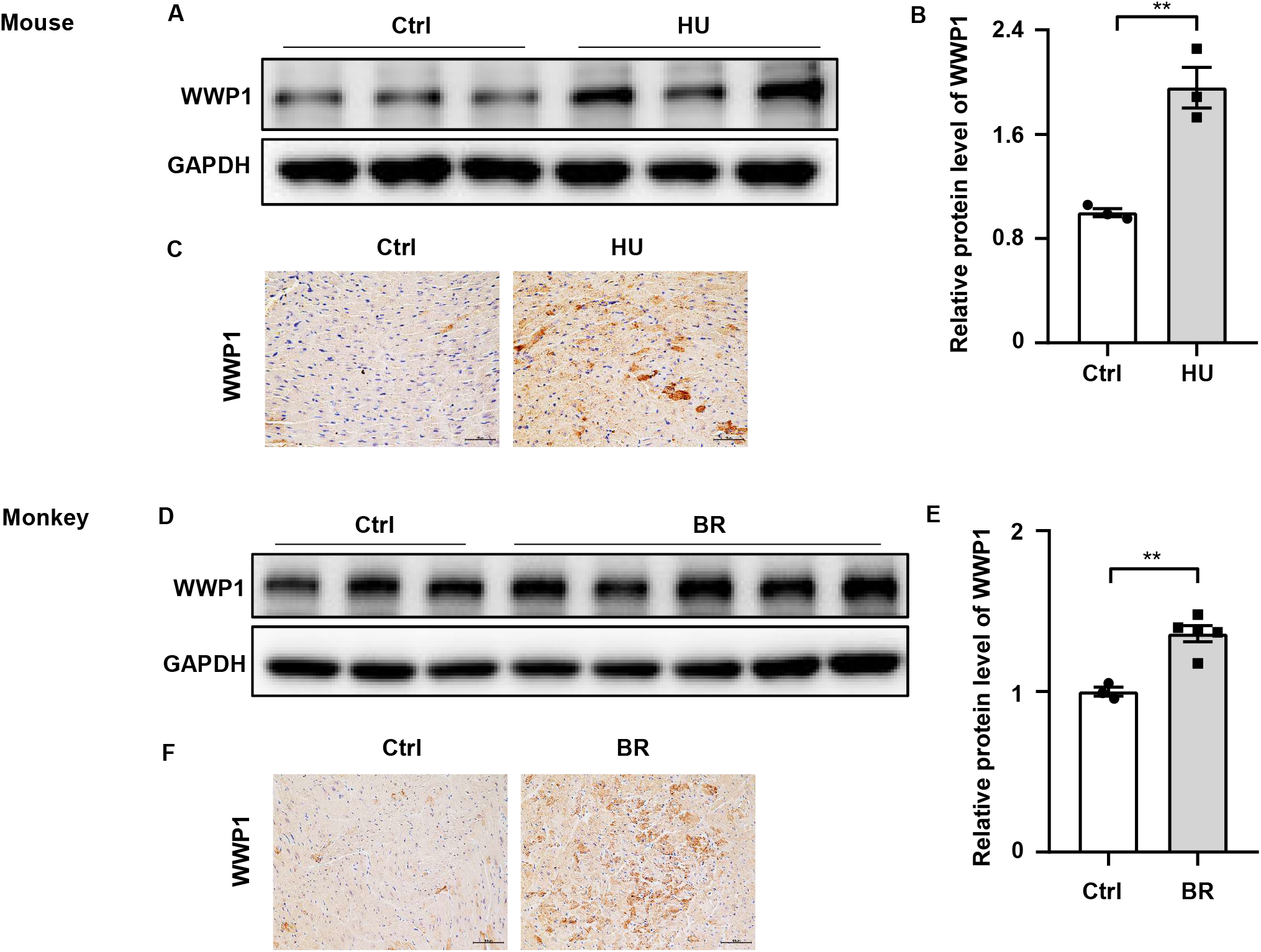
WWP1 expression changes in the hearts of mice and rhesus monkeys after simulated microgravity. **A**, Representative Western blotting analysis of WWP1 expression in cardiac extracts of adult mice following hindlimb unloading (HU) for 6 weeks (n=3 for each group). **B**, Quantification of WWP1 protein levels of **A**. **C**, Immunohistochemistry for WWP1 (brown) in paraffin section from mouse hearts at 6 weeks after hindlimb unloading. Scale bars, 50 μm. **D**, Representative Western blotting analysis of WWP1 expression in cardiac extracts of adult rhesus monkeys following bed rest (BR) for 6 weeks (Ctrl, n=3; BR, n=5). **B**, Quantification of WWP1 protein levels of **D**. **C**, Immunohistochemistry for WWP1 (brown) in paraffin section from rhesus monkey hearts at 6 weeks after bed rest. Scale bars, 50 μm. Data represent the means ± SEM. *P < 0.05, ** P < 0.01. WWP1, WW domain containing E3 ubiquitin protein ligase 1; HU, hindlimb unloading; BR, bed rest.

### WWP1 deficiency protects against simulated microgravity induced cardiac function declining

To further investigate the potential role of WWP1 in cardiac remodeling induced by simulated microgravity, we compared the responses of wildtype (WT) and WWP1 knockout (KO) mice to HU. WT and KO littermates at 3 months of age were subjected to hinlimb unloading by tail suspension for 6 weeks. The relevant control group were treated equally, except for the tail suspension. Body weight, heart weight (HW) and tibia length (TL) were recorded (**Figure 2C–E**), cardiac function was calculated by echocardiography (**Figure 2A-B**). All echocardiographic measurements were made while the heart rates of mice were maintained at 450–550 beats per minute. Compared with the control group, left ventricular ejection fraction (EF) and fractional shortening (FS) decreased significantly in WT mice after 6 weeks of hindlimb unloading, whereas no such changes in KO mice are apparent (**Figure 2A-B**), Moreover, the ration of heart weight to tibia length (HW/TL) decreased significantly in WT but not WWP1 KO mice after HU (**Figure 2F**), suggesting that WWP1 knockout protect against simulated microgravity induced cardiac atrophy and function declining.

**Figure 2.**
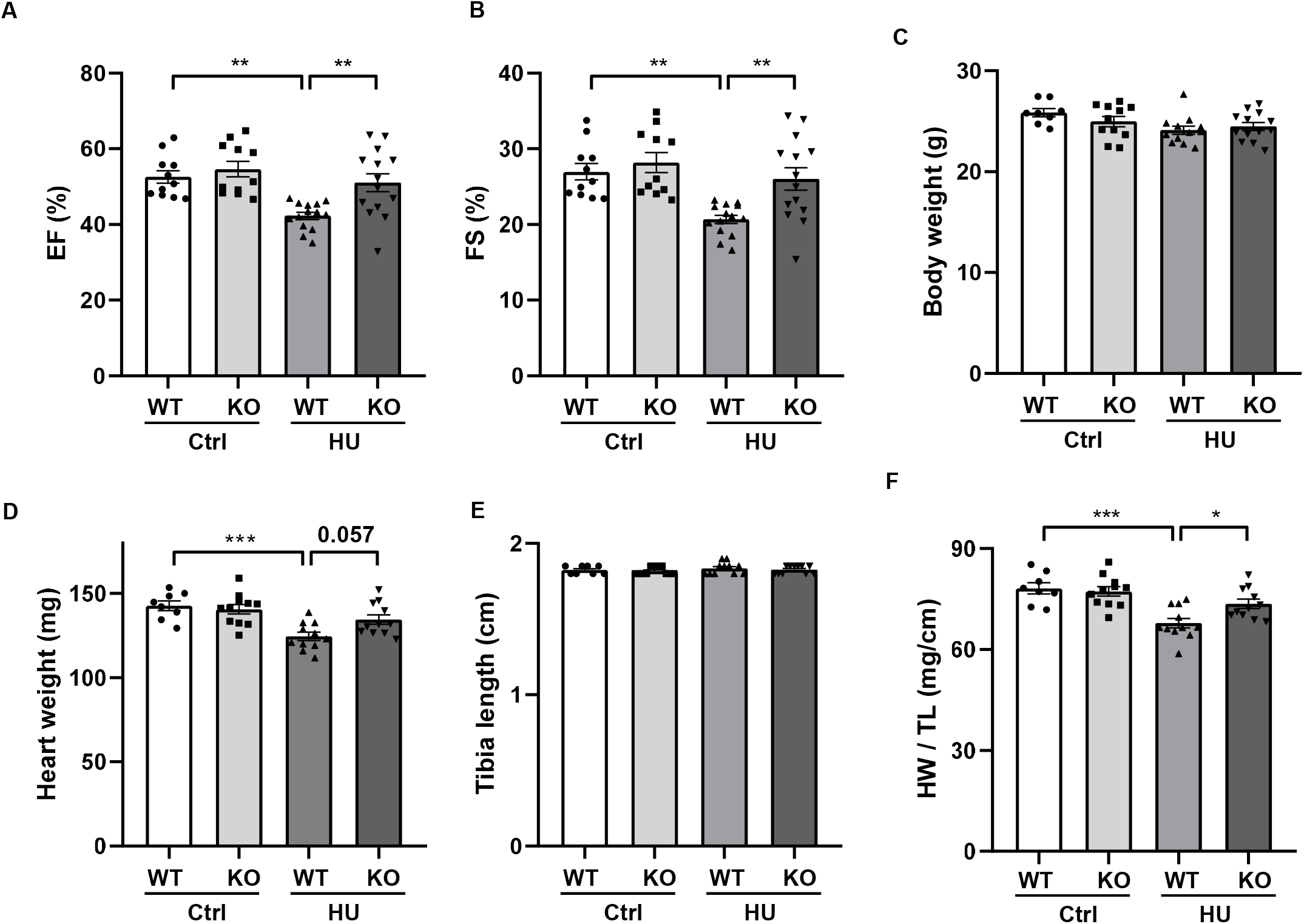
WWP1 knock out protects from simulated microgravity induced cardiac function declining. **A&B,** Echocardiographic assessment of ejection factor (EF) and fractional shortening (FS) of WT and WWP1 knockout mice after 6 weeks hindlimb unloading. **C-F**, Body weight, heart weight, tibia length, and the ratio of heart weight to tibia length of wild type (WT) and WWP1 knockout (KO) mice after 6 weeks hindlimb unloading. Data represent the means ± SEM. *P < 0.05, ** P < 0.01, ** *P < 0.001. Ctrl, control; HU, hindlimb unloading; WT, wild type mice, KO, knockout.

### Left ventricular structure of WT and KO mice following hindlimb unloading

To validate the influence of WWP1 KO in the heart structure after hindlimb unloading, we performed transthoracic echocardiography to determine the left ventricular structure of WT and KO mice that subjected to hindlimb unloading (**Figure 3**). A two way ANOVA was carried out on the end-diastolic left ventricular anterior wall thickness (LVAWs) by condition and genotype. There was a statistically significant interaction between the effects of condition and genotype on LVAWs. Compared with control, LVAWs of WT mice was decreased following hindlmb unloading, however LVAWs of WWP1 KO mice after hindlimb unloading did not change, and the value was higher than that in WT mice after hindlimb unloading (**Figure 3B**). The end-systolic left ventricular posterior wall thickness (LVPWs) showed the trend of decreasing in WT mice but not KO mice after hindlimb unloading, (**Figure 3D**). In the HU group, both the end-diastolic LV internal diameter (LVIDs) (**Figure 3F**) and the end-systolic LV volume (LV Vols) (**Figure 3H**) of KO mice are lower than that of WT mice. Meantime, the end diastolic anterior wall thickness (LVAWd) (**Figure 3A**), the end-diastolic left ventricular posterior wall thickness (LVPWd) (**Figure 3C**), the end-systolic LV internal diameter (LVIDd) (**Figure 3E**), and the end-systolic LV volume (LV Vold) (**Figure 3G**) did not change in the WT and KO mice after hindlimb unloading. These data indicated that WWP1 KO protected the heart from the decreasing of LVAWs induced by hindlimb unloading.

**Figure 3.**
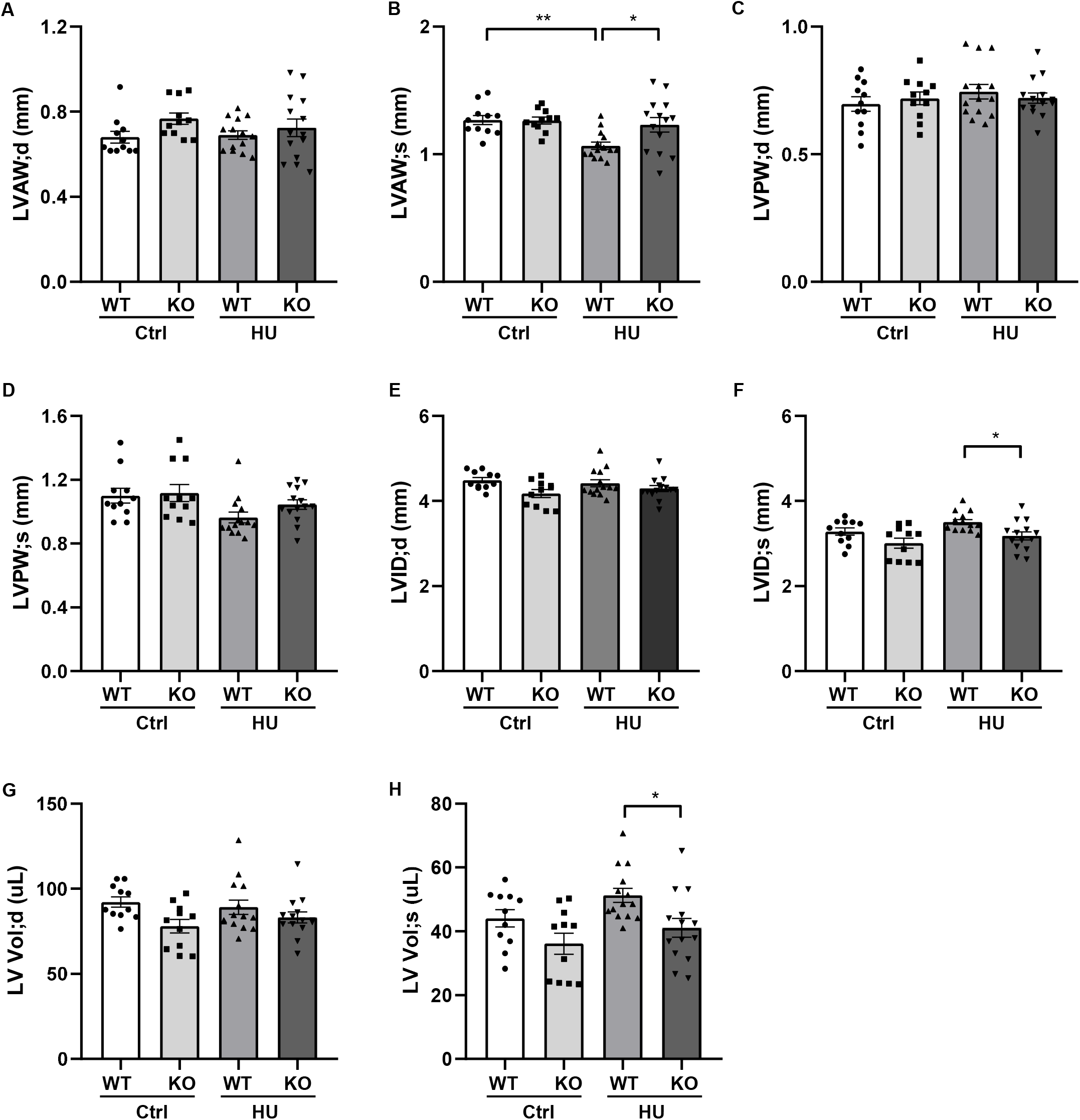
Transthoracic echocardiography evaluating the left ventricular structure of WT and WWP1 KO mice following hindlimb unloading. **A-H,** Quantitative analysis of the diastolic and systolic left ventricular posterior wall thickness (LVPWd and LVPWs), LV anterior wall thickness (LVAWd and LVAWs), LV internal diameter (LVIDd and LVIDs) and LV volume (LV Vold and LV Vols) from WT and WWP1 KO mice by echocardiography following hindlimb unloading. Data represent the means ± SEM. *P < 0.05, ** P < 0.01. Ctrl, control; HU, hindlimb unloading; WT, wild type mice, KO, knockout.

### WWP1 KO Protects against Simulated Microgravity Induced-Cardiac remodeling

To address the effect of WWP1 knockout on cardiac remodeling induced by hindlimb unloading, hearts from WT and KO mice were assessed for changes in morphology and cardiac remodeling genes expression. As shown in **Figure 4A**, Masson trichrome staining (MTT) showed a deeper staining of collagen in the heart of WT mice after hindlimb unloading, but WWP1 KO mice after hindlimb unloading had no obvious change. Wheat germ agglutinin (WGA) staining was used to demarcate cell boundaries of cardiomyocytes, compared with control, relative cell size was decreased in WT mice after hindlimb unloading, however, WWP1 KO can withstand the effect of hindlimb unloading (**Figure 4A-B**). The results of Q-PCR showed that transcripts for the pathological cardiac remodeling genes-BNP, Col1a1, and Col3a1 were significantly increased in the hearts of WT mice after hindlimb unloading, and WWP1 KO protected from the increase of cardiac remodeling genes expression induced by simulated microgravity (**Figure 4C-E**). These results suggested that WWP1 KO protects against simulated microgravity induced-cardiac remodeling.

**Figure 4.**
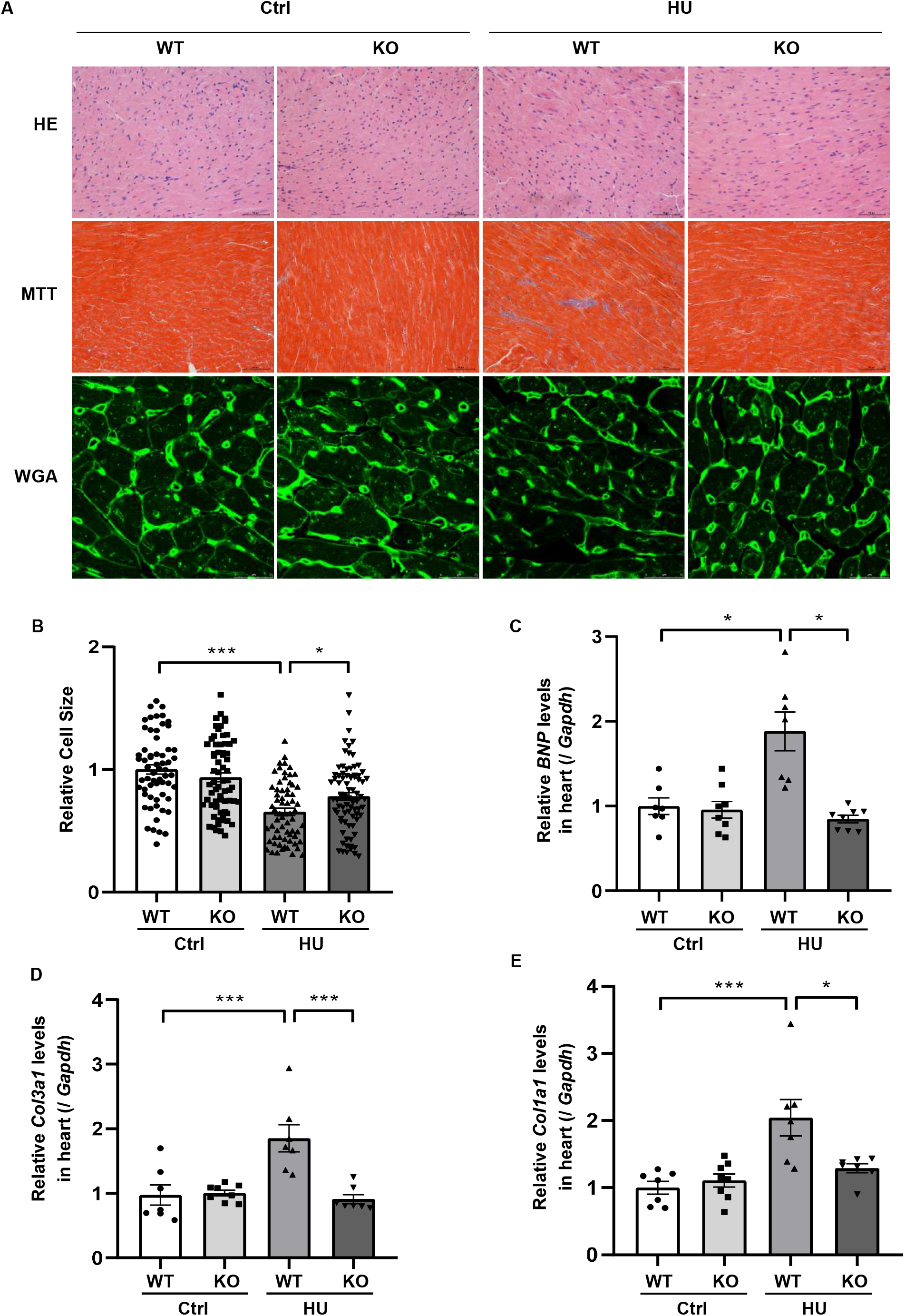
WWP1 knockout protects against simulated microgravity induced-cardiac atrophy. **A**, H&E-stained sections of hearts from WT and WWP1 KO mice after 6 weeks of hindlimb unloading. Sections of hearts are stained with Masson trichrome (MTT) to detect fibrosis (blue). Wheat germ agglutinin (WGA) staining is used to demarcate cell boundaries. Scale bars: 50 μm. **B**, The cardiomyocyte crosssectional area was measured from 7-μm-thick heart sections that had been stained with WGA by ImageJ software (NIH). Only myocytes that were round were included in the analysis. **C–E**, The mRNA levels of *BNP, Col1a1* and *Col3a1* were analyzed by Q-PCR from WT and WWP1 KO mice after 6 weeks of hindlimb unloading. The relative abundance of transcripts were quantified and normalized to *Gapdh*. Data represent the means ± SEM, *P < 0.05, **P < 0.01, **P < 0.001. *Col1a1*, alpha-1 type I collagen; *Col3a1*, alpha-1 type III collagen; BNP, brain natriuretic peptide; Q-PCR, real-time quantitative polymerase chain reaction.

### DVL2 positively correlates with WWP1 in heart of mice and rhesus monkeys after simulated microgravity

Our previous research found that WWP1 can participate in the regulation of myocardial remodeling by stabilizing DVL2^[9]^. In order to verify whether the above mechanism also exists in the myocardial remodeling caused by simulated microgravity, we analyzed the protein level of DVL2 in the heart of mice subjected to hindlimb unloading and rhesus monkeys subjected to bed rest. The protein level of DVL2 significantly increased (**Figure 5A-F**) and was significantly positive correlated with the expression of WWP1 in the heart of mice and rhesus monkeys subjected to simulated microgravity (**Figure 5G-H**), suggesting that WWP1 is correlated with DVL2 protein stability during simulated microgravity induced cardiac remodeling.

**Figure 5.**
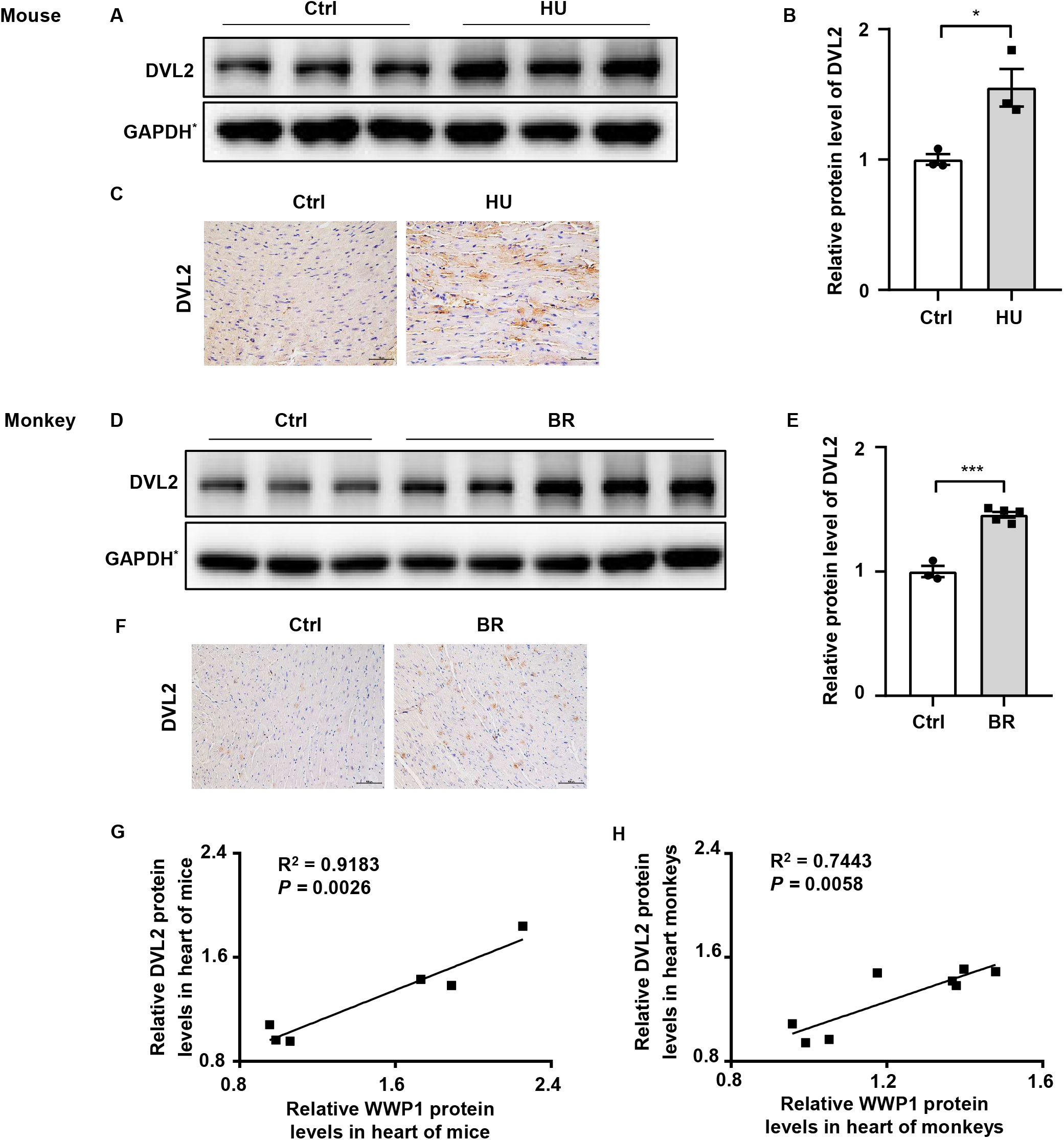
DVL2 expression changes in the hearts of mice and rhesus monkeys after simulated microgravity. **A**, Representative Western blotting analysis of DVL2 expression in cardiac extracts of adult mice following hindlimb unloading (HU) for 6 weeks (n=3 for each group). **B**, Quantification of DVL2 protein levels of **A**. **C**, Immunohistochemistry for DVL2 (brown) in paraffin section from mouse hearts at 6 weeks after hindlimb unloading. Scale bars, 50 μm. **D**, Representative Western blotting analysis of DVL2 expression in cardiac extracts of adult rhesus monkeys following bed rest (BR) for 6 weeks (Ctrl, n=3; BR, n=5). **E**, Quantification of DVL2 protein levels of **D**. **F**, Immunohistochemistry for DVL2 (brown) in paraffin section from rhesus monkey hearts at 6 weeks after bed rest. **G**, Pearson Correlation Coefficients between DVL2 and WWP1 protein levels in heart of mice following hindlimb unloading. **H**, Pearson Correlation Coefficients between DVL2 and WWP1 protein levels in heart of rhesus monkeys following bed rest. Scale bars, 50 μm. Data represent the means ± SEM. *P < 0.05, ***P < 0.001. DVL2, dishevelled segment polarity protein 2; HU, hindlimb unloading; BR, bed rest.

### WWP1 regulates simulated microgravity induced cardiac remodeling via DVL2/CaMKII/HDAC4 axis

To gain more insights into the effect of WWP1 KO on the signaling pathways involved in the cardiac remodeling induced by simulated microgravity, we we examined the protein level of DVL2, phosphorylation levels of CaMKII, and phosphorylation levels of HDAC4 in heart tissues of WT and KO mice after hindlimb unloading (**Figure 6A**). The two way ANOVA analysis reports showed there were statistically significant interactions between the effects of condition and genotype on the protein levels of DVL2, the phosphorylation levels of CaMKII at Thr287, and the phosphorylation levels of HDAC4 at Ser246. As shown in **Figure 6B-E**, quantifications analysis revealed that the level of DVL2, p-CaMKII and p-HDAC4 were increased in WT mice after hindlimb unloading, but it had no obvious changes in KO mice after hindlimb unloading. These results indicated that WWP1 regulates simulated microgravity induced cardiac remodeling via DVL2/CaMKII/HDAC4 axis.

**Figure 6.**
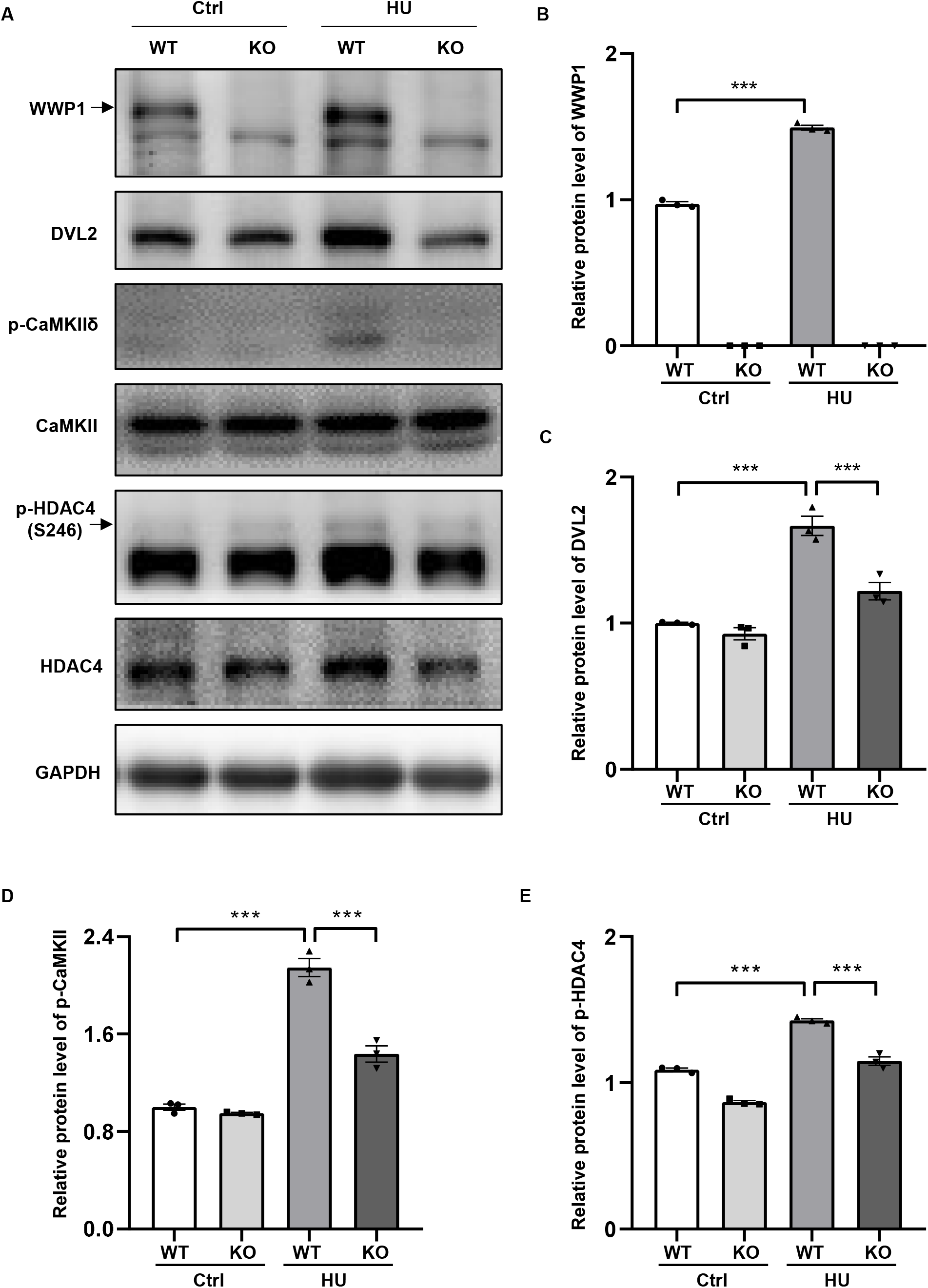
WWP1 knockout inhibits DVL2/CaMKII/HDAC4 signal in mice heart after simulated microgravity. **A,** Representative Western blots for WWP1, DVL2, CaMKIIδ and phosphorylation at (Thr287), HDAC4 and phosphorylation at Ser246 in hearts from WT and WWP1 KO mice after 6 weeks of hindlimb unloading. GAPDH levels served as a loading control. **B**, Quantification analysis of corresponding protein in **A**. Data represent the means ± SEM. ***P < 0.001. WWP1, WW domain containing E3 ubiquitin protein ligase 1; DVL2, dishevelled segment polarity protein 2; CaMKII, calcium/calmodulin-dependent protein kinase II; HDAC4, histone deacetylase 4; HU, hindlimb unloading.

## Discussion

Here, we identified WWP1 as a novel regulator of simulated microgravity-induced cardiac remodeling. WWP1 involved in the regulation of simulated microgravity-induced cardiac remodeling through DVL2/CaMKII/HDAC4/MEF2C pathway (**Figure 6**). WWP1 protein levels were significantly upregulated in the hearts of mice after 6 weeks of hindlimb unloading and rhesus monkeys after 6 weeks of bed rest, and the protein levels of WWP1 and DVL2 were closely related to the development of cardiac atrophy through the CaMKII/HDAC4-dependent pathway. WWP1 knockout protected against simulated microgravity induced-cardiac remodeling and function decline. Histological analysis demonstrated WWP1 KO inhibited the decrease in the size of individual cardiomyocytes of mice after hindlimb unloading. Moreover, the pathological cardiac remodeling signals, such as DVL2, CaMKII and HDAC4, were activated in the heart of WT mice after hindlimb unloading, however, WWP1 KO mice showed a different trend. Therefore, WWP1 represents a potential therapeutic target for cardiac remodeling and function decline induced by simulated microgravity.

The mammalian heart is a muscle, whose fundamental function is to pump blood throughout the circulatory system^[16]^. The cardiac muscle is well regulated in response to changes in loading conditions, typically caused by pathological or physiological stimulation^[4]^. A variety of stimuli can induce the heart to grow or shrink. Exercise, pregnancy, and postnatal growth promote physiologic growth of the heart^[6]^. With prolonged pressure overload, the heart undergoes pathologic hypertrophic remodeling, resulting in heart failure^[4]^. Cardiac atrophy was a complication for prolonged microgravity during space flight, long-term bed rest and mechanical unloading with a ventricular assist device^[3, 4, 17, 18]^. The tail suspension (hindlimb unloading) of mice or rats is widely utilized to study the effects of microgravity^[19]^, and head-down tilt bed rest model for non-human primate-rhesus monkeys or human volunteers is also a classical ground-based model of microgravity^[3, 20, 21]^. Our early researches also demonstrated that simulated microgravity induced cardiac atrophy and function decline of mice and rhesus monkeys^[3, 6]^. In this study, we found that hindlimb unloading of mice led to cardiac atrophy, as evidenced by decreased LVAWs, HW/TL and relative cell size, and WWP1 deficiency can effectively combat this situation.

Although myocardial hypertrophy and myocardial atrophy have opposite phenotypes, they show similar gene expression characteristics, such as “fetal” gene expression activation, supporting the concept that opposite changes in workload in vivo induce a similar transcriptional response^[22]^. Compared with other forms of cardiac remodeling, little is known about the specific mechanism governing the microgravity-induced cardiac atrophy^[3]^. To systematically explore the effects of spaceflight on the heart, Walls et al. used the Drosophila cardiac model to examine how prolonged exposure to microgravity affects cardiac health. They reported the effects of microgravity on heart function in wild-type (WT) fruit flies that were born, developed, and spent 1–3 weeks as adults aboard the ISS compared with their ground-based controls, the results showed that structural and functional cardiac remodeling occurs in response to microgravity, RNA sequencing and immunohistochemistry data suggested that alterations in proteostasis likely contribute to the observed changes in cardiac remodeling induced by microgravity^[1]^. According to their research, cardiac atrophy mediated by protein homeostasis regulation may be a fundamental response of heart muscle to microgravity^[1]^. Protein ubiquitination is a multi-functional post-translational modification that affects a variety of diseases processes, including cardiac remodeling^[23]^. Our previous research found that E3 ubiquitin ligase WWP1 can stabilize DVL2 by catalyzing K27-linked polyubiquitination, DVL2 positively correlated with WWP1 hypertrophic heart and mediated the regulation by WWP1 of the CaMKII/HDAC4/MEF2C axis in cardiac hypertrophy^[9]^. In this study, we found that the protein levels of WWP1 and DVL2 were significantly upregulated in the hearts of mice and rhesus monkeys after simulated microgravity, and WWP1 deficiency can protect against simulated microgravity induced-cardiac remodeling by inhibition of DVL2/CaMKII/HDAC4 pathway.

Well-characterized signaling molecules that regulate cardiac remodeling induced by pressure overload include DVL2 (Zhao et al., 2021), Calcium/calmodulin-dependent kinases (CaMKII)^[24]^ and histone deacetylase 4 (HDAC4)^[8]^. DVL2 is highly evolutionarily conserved and participate in canonical and non-canonical Wnt pathways^[25]^. In particular, DVL2 acts as an activator in pressure overload induced cardiac hypertrophy^[9, 26]^. CaMKII, an important regulator of non-canonical Wnt signaling, whose continuously activated form is critical in pathological cardiac remodeling^[24, 27]^. The activation of CaMKII directly phosphorylates HDAC4, causing its relocalization to the cytoplasm and activation of myocyte enhancer factor 2C (MEF2C)^[8, 28, 29]^. The protein level of DVL2 and the phosphorylation level of CaMKII was increased in the heart of WT mice, but not WWP1 KO mice, subjected to hindlimb unloading (**Figure 6A**). The phosphorylation level of HDAC4 was similar to that of CaMKII (**Figure 6A**). In summary, we have discovered that WWP1 is involved in the regulation of simulated microgravity induced cardiac remodeling through DVL2/CaMKII/HDAC4/MEF2C pathway, moreover, WWP1 has potential as a therapeutic target for cardiac remodeling induced by simulated microgravity.

## Conflict of Interest

The authors report no conflicts of interest whether financial or otherwise.

## Author Contributions

Conceptualization: YXL and SKL; Methodology: GHZ, ZFL and XXY; Validation: GHZ; Formal Analysis: GHZ and SKL; Investigation: GHZ, DSZ, JWL, ZFL, JJP and WJX; Writing-Original Draft: GHZ; Writing—Review and Editing: GHZ, YXL and SKL; Funding Acquisition: GHZ, YXL and SKL.

## Funding

This work was supported by the National Natural Science Foundation of China (No. 81822026, 81701859, 31670865, and 81830061), the Grant of State Key Lab of Space Medicine Fundamentals and Application (SMFA17B05 and SMFA19A02), Space Medical Experiment Project of China Manned Space Program (NO. HYZHXM01007).

## Notes

### Competing Interest Statement

The authors have declared no competing interest.

